# Failed Cellular Surveillance Enables Pathogenic Matrix Deposition in a *COL2A1*-Related Osteoarthritis

**DOI:** 10.1101/2023.10.19.562780

**Authors:** Kathryn M. Yammine, Sophia Mirda Abularach, Michael Xiong, Seo-yeon Kim, Agata A. Bikovtseva, Vincent L. Butty, Richard P. Schiavoni, John F. Bateman, Shireen R. Lamandé, Matthew D. Shoulders

## Abstract

Mutations in the *COL2A1* gene, encoding procollagen-II, cause various chondrodysplasias, including precocious osteoarthritis with mild spondyloepiphyseal dysplasia engendered by the p.Arg719Cys substitution. The molecular mechanisms underlying these disorders remain incompletely understood, largely owing to the absence of models faithfully recapitulating the human disease. Here, we developed an *in vitro* human cartilage model using isogenic induced pluripotent stem cell (iPSC) lines carrying either wild-type or Arg719Cys *COL2A1*. Directed differentiation into chondrocytes yielded cartilage tissues that were analyzed by immunohistochemistry, electron microscopy, SDS-PAGE, and RNA-sequencing. Tissues derived from Arg719Cys heterozygotes displayed a deficient matrix, closely reflecting the human disease phenotype. Arg719Cys procollagen-II was excessively post-translationally modified and partially retained within the endoplasmic reticulum (ER), leading to ER distention. Notably, despite introduction of an aberrant cysteine residue—expected to engage redox-sensitive folding and quality control pathways—Arg719Cys procollagen-II was not detectably recognized by the ER proteostasis network. The resulting inability to mount a quality control response, including activation of the unfolded protein response, indicates a failure in cellular surveillance. As a result, malformed procollagen-II both accumulates intracellularly and is secreted, contributing to the deposition of a structurally compromised extracellular matrix that drives disease pathology. The iPSC-derived cartilage model presented here provides a genetically defined and expandable, human-based system for dissecting the mechanisms of failed proteostasis in collagenopathies. These findings shed light on the types of substitutions in procollagen that cells can or cannot recognize, and underscore the therapeutic potential of targeting cellular surveillance and collagen quality control pathways in *COL2A1*-related disorders and beyond.

**SIGNIFICANCE:** The p.Arg719Cys substitution in *COL2A1* is known to cause dominantly inherited precocious osteoarthritis with mild spondyloepiphyseal dysplasia, but its molecular pathogenesis remains poorly understood. Prior work suggests disrupted fibrillogenesis as the molecular etiology, yet phenotypically relevant human models are lacking. Here, we use human iPSC-derived cartilage to uncover how the Arg719Cys substitution alters procollagen-II folding, modification, trafficking, and matrix deposition. We observe hallmark disease phenotypes, including a sparse collagen-II matrix and distended endoplasmic reticulum (ER), consistent with intracellular accumulation of procollagen-II. Surprisingly, the mutant procollagen-II does not appear to be differentially engaged by the ER proteostasis network, despite the incorporation of an aberrant and apparently non-disulfide bond-forming Cys residue, suggesting a fundamental failure of cellular surveillance. This unrecognized misfolding leads to secretion of defective collagen culminating in matrix dysfunction. These findings establish a mechanistic basis for disease etiology and highlight collagen quality control as a potentially tractable therapeutic target. The iPSC-based model provides a scalable, human-relevant platform for dissecting disease mechanisms and developing therapies for *COL2A1*-related disorders and other collagenopathies.

## INTRODUCTION

Mutations in the gene encoding procollagen-II (*COL2A1*) cause a variety of disorders affecting joint health and skeletal development, termed type-II collagenopathies.^1–3^ The vast majority of these diseases are caused by autosomal dominant mutations in the triple-helical region of procollagen-II.^4^ Mutations leading to Arg➔Cys substitutions are a common subtype of collagenopathy-inducing mutations.^5^ A prominent example is the *COL2A1* p.Arg719Cys substitution (exon 33 c.2155C>T; previously Arg519Cys) substitution, which has been identified in several families from at least three distinct founders.^6–8^ Affected individuals present with mild spondyloepiphyseal dysplasia and precocious generalized osteoarthritis.^7^ Currently, there is no cure. Treatment often involves multiple surgical joint replacements.^9^

A mechanistic understanding of the collagenopathies is still lacking. Much work in the field has focused on Gly substitutions,^10, 11^ which disrupt triple-helix folding by eliminating the essential Gly residue at every third position in the triple helix.^2^ A recent report from our group described the endoplasmic reticulum (ER) storage defect created in a human cartilage model by the Gly1170Ser procollagen-II substitution.^12^ In that genotype and system, the ER proteostasis network robustly recognized the slow-folding procollagen that accumulated in the ER, consistent with the need to re-initiate stalled triple-helix folding downstream of the Gly substitution.^12^

While less common than Gly substitutions, Arg➔Cys substitutions in the triple-helical domain are among the most common disease-causing mutations in fibrillar collagens.^5^ The mechanism by which Arg➔Cys substitutions might cause collagenopathies is unclear, as disruption of triple-helix folding is not an obvious consequence of such a substitution.^13, 14^ One possibility is that the introduction of an aberrant Cys residue, capable of forming covalent bonds with diverse proteins in the oxidative ER environment, stimulates inappropriate disulfide bond formation and/or extensive interactions with the ER’s redox-related proteostasis machinery.

Another possibility is that loss of the Arg residue is itself problematic. However, to date, only Arg719Cys is reported to be pathogenic despite substitutions of Arg719 to all of Leu, Gly, His, and Cys being observed at low frequencies within the population.^15^ Further support for the idea that it is not loss of the Arg itself which is problematic is that this portion of the triple-helix is not known to lie in a recognition site for protein partners to interact with collagen-II in the extracellular matrix, as previously assessed by binding assays.^11, 16, 17^

Efforts to uncover the molecular mechanism of disease have been hampered by the need for a relevant disease model of cartilage, the main tissue affected by the Arg719Cys substitution in procollagen-II. Primary patient cells, beyond being scarce and difficult to procure outside of a joint replacement, are challenging to study *in vitro* owing in large part to an inability to culture primary chondrocytes without dedifferentiation.^18^ Studies on patient-derived tissue suggest that Arg719Cys procollagen-II engages in an abnormal disulfide bonding pattern in the matrix,^19^ a notable finding given that recombinant Arg719Cys procollagen-II studied *in vitro* does not,^20^ further emphasizing the need for a robust model system. Two related transgenic mouse models of this substitution have been developed, showing that mice homozygous for the human Arg719Cys procollagen-II transgene have disrupted growth plate organization and reduced collagen-II fibril density.^21, 22^ While useful, results from these models must be cautiously interpreted as they involve the expression of a mutated human *COL2A1* gene in a murine background. Overall, there continues to be a need for a model that accurately reflects the genetics of the disease in a relevant human cellular context, both to advance our understanding of the etiology of this collagenopathy and to enable the discovery of potential treatments.

Here, enabled by directed differentiation of induced pluripotent stem cells (iPSCs),^12, 23, 24^ we introduce a human cartilage model of precocious osteoarthritis with mild spondyloepiphyseal dysplasia and assess the effects of the heterozygous Arg719Cys substitution in procollagen-II. Critically, our human iPSC-based model of the pathologic Arg719Cys substitution is paired with an isogenic, wild-type control,^25^ allowing for direct phenotypic evaluation of the pathologic substitution without confounding genetic effects. Further experiments reveal, unexpectedly, that the ER proteostasis network does not differentially engage this disease-causing, redox-active residue-substituted procollagen-II – a failure that then propagates to disease-causing matrix defects. Beyond the mechanistic insights obtained on the types of procollagen mutations capable of evading cellular detection, the scalable human cartilage model introduced here provides a unique and phenotypically robust platform to, for the first time, develop and test therapeutic strategies targeting procollagen folding and secretion in collagenopathies driven by non-Gly substitutions in the collagen triple-helical domain.

## RESULTS

### Arg719Cys procollagen-II-expressing cells can be differentiated into chondrocytes that robustly deposit cartilage extracellular matrix

Using an established protocol,^23^ we differentiated previously validated,^25^ genetically matched wild-type and Arg719Cys iPSC lines into chondrocytes. Upon differentiation to chondrocytes, both lines deposited cartilaginous extracellular matrices rich in collagen-II (**Fig. 1A**) and proteoglycans (**Fig. 1B**). Immunohistochemistry revealed no evident differences in the collagen-II or proteoglycan staining between genotypes. To further evaluate successful differentiation to chondrocytes and more broadly compare possible matrix composition defects, we examined expression of matrisome genes^26^ by RNA-sequencing. Consistent with successful chondrocyte differentiation, cartilage-specific genes including *COL2A1, COL9A1, COL11A1* and *ACAN* were the most highly expressed matrisome transcripts in both lines (**Fig. 1C**). Matrix genes specific to other tissue types were not substantially expressed, including *COL1A1* – an abundant matrix protein in bone and other tissues – confirming successful differentiation to chondrocytes. Moreover, comparison of gene expression across genotypes revealed only minor changes in matrisome expression associated with the pathologic Arg719Cys procollagen-II (**Fig. 1D**). Just six matrisome genes were differentially expressed >2-fold, consistent with the matrix composition being largely similar between genotypes. In sum, expression of Arg719Cys substituted procollagen-II prevented neither chondrogenesis nor bulk matrix deposition.

**Fig. 1.**
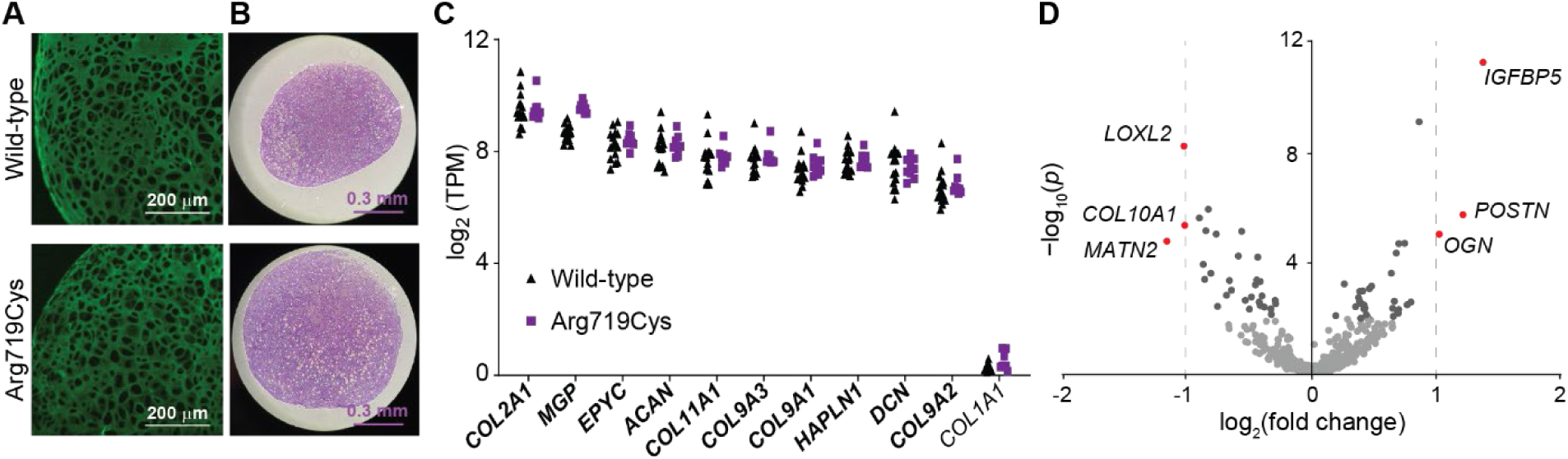
**(A)** Immunohistochemistry against collagen-II revealed that both the wild-type and heterozygous Arg719Cys procollagen-II matrices were rich in collagen-II. Scale bar represents 200 μm. **(B)** Staining histological sections with toluidine blue indicated that both matrices were also similarly rich in proteoglycans. Scale bar represents 0.3 mm. **(C)** Cartilage-specific genes were the most highly expressed of the matrisome genes in both wild-type and Arg719Cys procollagen-II expressing cells, as evidenced by RNA-sequencing at day 44. Genes associated with other tissues (e.g., *COL1A1*) were not substantially expressed. TPM: transcripts per million. At least nine individual chondronoids per genotype were analyzed. **(D)** Differential expression analysis of the matrisome at day 44 revealed only a few genes significantly differentially expressed between genotypes. Fold-change is presented here as the gene expression in Arg719Cys procollagen-II-expressing cells compared to wild-type. Genes with a significance of −log_10_(*p*) > 2 are depicted in dark grey, while those more than 2-fold differentially expressed [log_-_(fold change) > 2 or <−2] are depicted in red. Significance was evaluated using the Wald test on read count data modeled in a negative binomial framework (DESeq2 algorithm).

### Arg719Cys procollagen-II leads to a deficient cartilage matrix

Given its association with disease, we anticipated that the Arg719Cys substitution would cause deposition of a defective collagen-II matrix. Hence, despite its apparently normal appearance when evaluated via low magnification immunohistochemistry (**Fig. 1A**), we still sought to examine the integrity of the collagen-II matrix more closely using transmission electron microscopy (TEM). Whereas the wild-type matrix was rich in fibrils, we observed that the matrix of heterozygous cells expressing Arg719Cys procollagen-II was deficient by TEM. Representative images of the matrix of both genotypes are presented in **Fig. 2**. In particular, collagen fibrils were notably sparser in the disease variant, indicating that the Arg719Cys substitution leads to deposition of a defective cartilage collagen-II matrix.

**Fig. 2.**
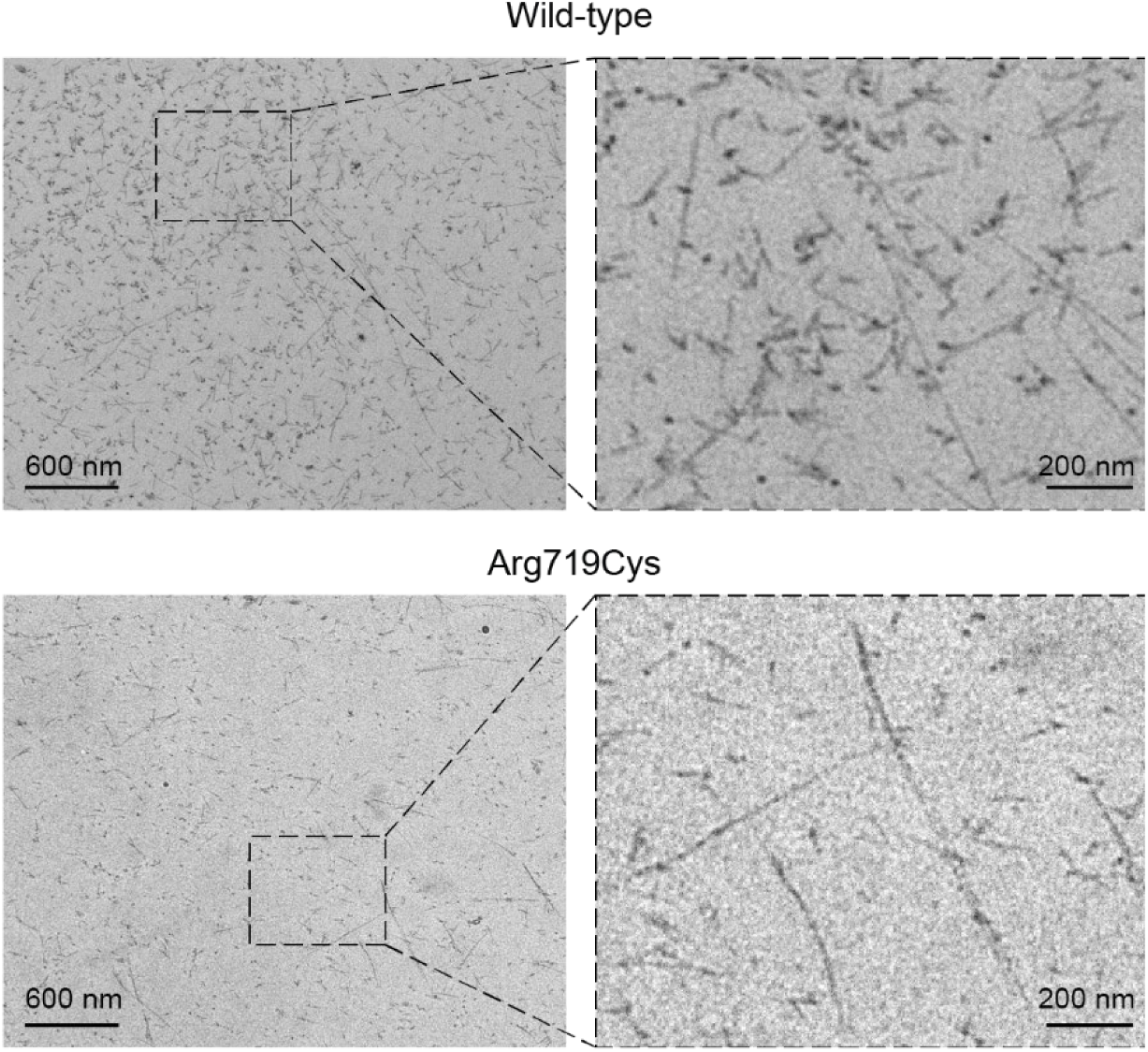
Compared to wild-type procollagen-II expressing chondronoids, transmission electron microscopy imaging of the matrix revealed a sparse network of collagen fibrils deposited by cells expressing the disease-causing procollagen-II variant Arg719Cys at day 44. Scale bars represent 600 nm and 200 nm, respectively.

### Arg719Cys procollagen-II is unexpectedly slow to fold and may accumulate intracellularly

Disease-causing Gly➔Ser substitutions in procollagen’s triple-helical domain commonly result in slowed triple-helix folding,^12, 27–31^ a phenomenon typically attributed to disruption of the essential network of Gly-requiring hydrogen bonds that stitch together the three distinctive strands of the triple helix.^2^ Arg➔Cys substitutions are not expected to disrupt this network, and so slowed triple-helix folding is not a presumptive outcome.^14, 20^ Relatedly, Arg➔Cys substitutions have been shown to have variable effects on the thermal stability of collagen-II based on their location within the triple helix;^14, 32^ Arg719Cys in particular was found not to decrease the stability of the triple helix.^20^ Still, given the defective cartilage-like matrix we observed, to evaluate this possibility we employed a gel-based analysis widely used in the field to assess the rate of triple-helix formation in cells. Specifically, as procollagen folds in the ER, its unfolded regions accumulate post-translational modifications, especially hydroxylated lysine residues that are then *O*-glycosylated.^33^ Once the triple-helix forms, modification is halted. Thus, if a mutation slows triple-helix formation, the procollagen accumulates excess *O*-glycans, which are identifiable by a reduction in electrophoretic mobility on SDS-PAGE.^27–30, 34^

We extracted collagen from the iPSC-derived cartilage tissue and separated the protein using SDS-PAGE. We observed a modest reduction in Arg719Cys-substituted collagen-II’s electrophoretic mobility when compared to wild-type collagen-II (**Figs. 3A** and **S1**), indicating that the Arg719Cys substitution does indeed slow the rate of triple-helix formation in cells.

Notably, this finding recapitulates the slow migration phenotype of Arg719Cys collagen-II extracted from patient tissue, further confirming the relevance of our model to the human disease.^19^

We next turned to microscopy to evaluate the fate of slow-folding Arg719Cys procollagen-II. Using immunohistochemistry against procollagen-II, we noted that more procollagen-II is retained intracellularly in cells expressing the disease variant (**Fig. 3B**). For example, blinded quantification revealed that, on average, 51% of cells expressing Arg719Cys substituted collagen-II contained intracellular procollagen-II, compared to only 34% of wild-type cells (**Fig. 3C**). Co-staining with an ER marker revealed that the majority of the intracellular procollagen-II was retained in the ER. Moreover, when examined by TEM, we observed that cells expressing the disease variant generally exhibited dilated ER (**Fig. 3D**), further supporting the notion that the slow-folding Arg719Cys procollagen-II accumulates to some extent in the ER.

**Fig. 3.**
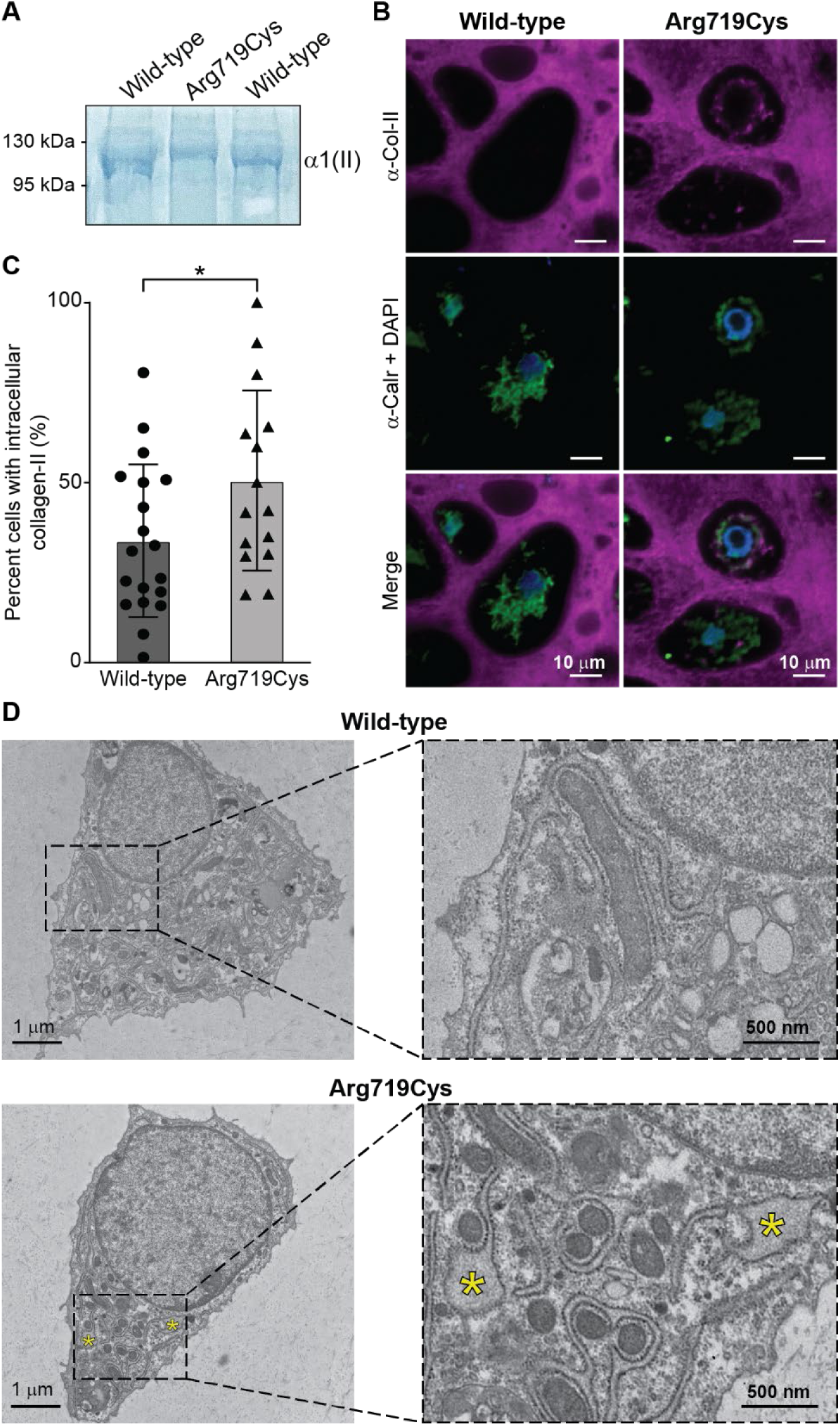
**(A)** SDS-PAGE of collagen extracted from cartilage produced by wild-type or Arg719Cys procollagen-II expressing cells at day 44. Arg719Cys collagen-II exhibited a reduction in electrophoretic mobility compared to wild-type. **(B)** Confocal imaging of chondronoid sections immunostained against collagen-II and the ER marker calreticulin (Calr) at day 44 revealed intracellularly retained procollagen-II in the disease variant. Scale bars represent 10 μm. **(C)** Blinded quantification suggested that Arg719Cys procollagen-II-expressing cells may contain more intracellular procollagen-II, as compared to wild-type. * *p* < 0.05 (*p* = 0.043), as evaluated using an unpaired, two-tailed *t* test. Each data point represents a single confocal image. Error bars represent standard deviation. **(D)** Expression of Arg719Cys procollagen-II was associated with distended ER (marked with yellow asterisks), as evidenced by TEM at day 44. Scale bars represent 1 μm and 500 nm, respectively.

We emphasize that both the intracellular retention and dilated ER phenotypes are relatively mild compared to other pathologic substitutions – including the predicted to be more severe Gly substitutions.^12^ Nonetheless, these observations correlate with the relatively mild phenotype of this collagenopathy (normal development and habitus^6, 7^) on the spectrum of chondrodysplasias, which, at their most severe, are perinatal lethal.^35^

### Arg719Cys substituted procollagen-II is not recognized by the ER proteostasis network

Biochemically, introduction of a Cys in a domain of procollagen typically devoid of Cys could be very problematic, owing to the ability of the amino acid’s side chain to engage in unwanted disulfide bonds. Notably, the introduction of such an aberrant Cys residue opens additional opportunities for redox-active components of the ER proteostasis network, such as protein disulfide isomerases and other reductases, to engage the slow-folding protein. We therefore anticipated that Arg719Cys procollagen-II would be robustly engaged by the ER’s protein folding and quality control machinery.

To test this hypothesis, we used quantitative mass spectrometry to elucidate and compare the interactomes^12, 36–38^ of wild-type versus Arg719Cys-substituted procollagen-II under oxidizing conditions (**Fig. S2**). Owing to the lack of immunoprecipitation-quality procollagen-II antibodies to enable this experiment in the more biologically relevant chondronoids, we performed this study using overexpression of epitope-tagged procollagen-II constructs in HT-1080 cells, which other work has shown are capable of properly handling fibrillar collagens, but that do not express meaningful amounts of endogenous interfering fibrillar collagen.^36, 39^

Of the identified high-confidence procollagen-II interacting proteins, we first focused on those capable of engaging in a disulfide bond with the pathologic Cys719, including protein disulfide isomerases and dithiol-disulfide exchange enzymes. None of the six disulfide bonding-capable proteins detected interacted differentially with Arg719Cys procollagen-II compared to wild-type (**Table 1**), suggesting that it is not covalent interaction with an unwanted partner during procollagen folding in the ER that slows folding and underlies pathology. Even more surprising, across all the >40 high confidence procollagen-II interacting proteins identified, none of the interactors – disulfide-forming or otherwise – significantly differentially engaged Arg719Cys procollagen-II compared to wild-type (**Table S1**). Thus, bearing in mind the limits of sensitivity in our analysis (which nonetheless covered the majority of known procollagen interactors), it appears that Arg719Cys procollagen-II is not differentially recognized by the ER proteostasis network (including chaperones, modifying enzymes, and proteins involved in quality control and degradation), at least in HT-1080 cells. Such a failure, in striking contrast to the Gly1170Ser-substituted procollagen-II that is robustly recognized by the ER proteostasis network,^12^ likely underlies the deposition of a defective matrix and disease.

### Consistent with failed recognition by the ER proteostasis network, expression of Arg719Cys-substituted procollagen-II does not induce a global stress response

Accumulation of misfolding proteins in the ER is often associated with activation of the unfolded protein response (UPR), which maintains proteostasis in the secretory pathway.^40^ On the other hand, the unchanged extent of interaction with BiP (**Table S1**), which controls UPR activation, suggested that such activation might not occur.^40, 41^ Failure to mount a beneficial repair response could be damaging. On the other hand, chronic stress could also drive disease.^31^ To untangle the relative contribution of these two phenomena, we turned to RNA-sequencing to elucidate the cellular processes at play in the phenotypically relevant chondronoid.

We assessed activation of the UPR, and various other possibly activated stress pathways, in Arg719Cys-expressing chondrocytes versus wild-type at two different timepoints selected to avoid overlap with chondrogenesis, a process known to involve a highly coordinated, but transient, UPR:^42^ day 34 and day 44 (**Fig. 4**). We observed that the UPR was not globally induced in the Arg719Cys cells, at either timepoint (**Fig. 4A**). Only one UPR-related gene was significantly upregulated, and exclusively at the earlier timepoint. Results were similar for other cellular stress responses, including a previously described non-canonical stress response to procollagen-I misfolding (**Fig. 4B**),^43^ the integrated stress response (**Fig. 4C**),^44^ inflammation (**Fig. 4D**), and oxidative stress (**Fig. 4E**). Despite modest changes in a few genes, we did not observe significant global differences in expression levels in the Arg719Cys-expressing cells compared to wild-type.

**Fig. 4.**
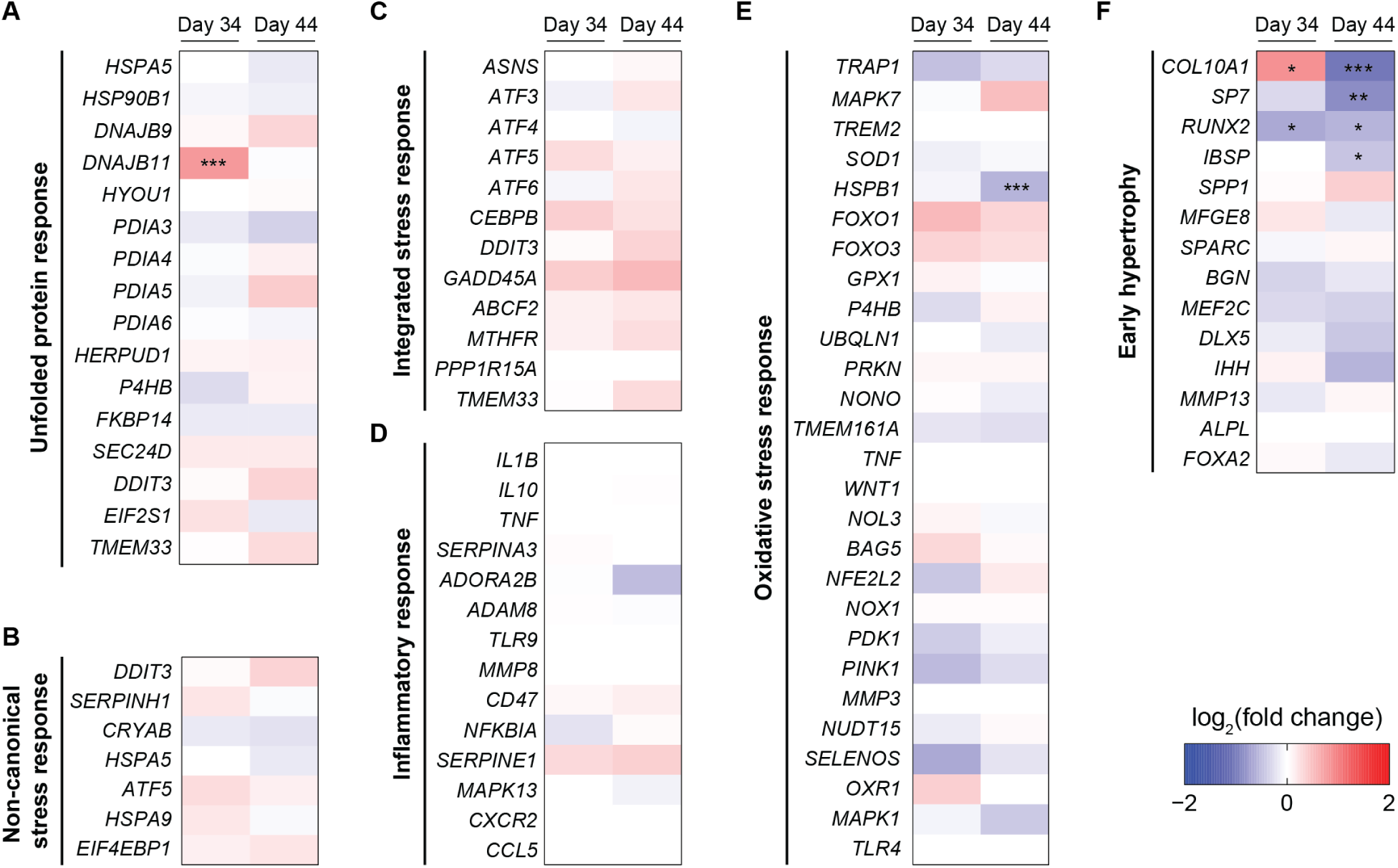
**(A–F)** Heatmap of the fold change gene expression in of Arg719Cys procollagen-II-expressing cells compared to wild-type at two timepoints. While a few individual genes were differentially expressed, no stress response was globally activated at the transcriptional level. * *p* < 0.05; ** *p* < 0.01; *** *p* < 0.001. Significance was evaluated using the Wald test on read count data modeled in a negative binomial framework (DESeq2 algorithm).

Gene set enrichment analyses (**Fig. S3**) similarly did not identify any differentially activated stress responses, although (exclusively at the early timepoint) they did highlight subtle differences in a few cellular processes, most notably translation. Overall, the RNA-sequencing data revealed that expression of Arg719Cys was not associated with global induction of a stress response at the transcriptional level in chondrocytes. Consistent with this finding, we did not observe a significant increase in apoptosis in cells expressing Arg719Cys, as assessed by TUNEL assay (**Fig. S4**). We did, however, note that, at the late timepoint (day 44), genes associated with early hypertrophy^23^ (a critical step in the endochondral ossification process) were significantly downregulated in cells expressing the disease variant (**Fig. 4F**).

These results further confirm that the cell is failing to recognize slow-folding, disease-causing Arg719Cys-substituted procollagen-II. That failure propagates to cartilage defects that then drive pathologic symptoms in joints.

## DISCUSSION

Herein, we characterized pathologic consequences of the autosomal dominant p.Arg719Cys mutation in *COL2A1* by developing a novel human iPSC-based cartilage model of the disease. Thanks to the genetically matched wild-type control,^25^ we were able to rigorously define the subtle defects that arise specifically owing to expression of this pathologic procollagen-II variant. We found that, despite their ability to deposit a cartilaginous matrix, heterozygous cells expressing Arg719Cys-substituted procollagen-II did display signs of pathology, including deposition of a structurally deficient collagen-II matrix, modest intracellular accumulation of procollagen-II, and ER distention. The relatively mild phenotype observed in this Arg719Cys procollagen-II cartilage model appears to be consistent with the severity of this disease on the chondrodysplasia spectrum.^7^ Moreover, the phenotypes described in this first human cartilage model of the disease are consistent with previous reports, including the observation of slow folding and hyper-modification of collagen-II extracted from patient cartilage,^19^ the disruption of fibril formation *in vitro*^20^ and *in vivo,*^21, 22^ and even the delay in early hypertrophy.^21, 22^ Overall, this human iPSC-based cartilage model appears to accurately recapitulate the genetics and phenotypes of Arg719Cys procollagen-II pathology, rendering it a powerful system for mechanistic and therapeutic discovery.

While the sparse collagen matrix could be a result of reduced collagen-II secretion, it may also be associated with dysregulated fibril formation. *In vitro* studies have shown that homotrimers of Arg719Cys collagen-II do not assemble into fibrils, and their presence in a mixture of wild-type collagen-II alters the self-assembly time and morphology of fibrils.^20, 45^ While heterotrimers of wild-type and Arg719Cys procollagen-II are statistically far more likely than Arg719Cys homotrimers when expressed in disease-relevant cells heterozygote for the mutation, like those used here, the excess post-translational modifications from the slow folding of the triple helix could still contribute to defects in fibril formation and crosslinking. Lower expression of lysyl oxidase-like 2 (LOXL2), an enzyme that catalyzes the first step in formation of collagen crosslinks,^46^ in chondrocytes expressing Arg719Cys procollagen-II (**Fig. 1D**) further supports the idea of a defect in crosslinking. POSTN (**Fig. 1D**) has also been suggested to impact lysyl oxidase function through matrix remodeling.^47, 48^

Introduction of a Cys residue in the triple-helical domain of procollagen-II might be predicted to be very problematic for procollagen folding. The aberrant Cys could engage in unwanted disulfide bonds with procollagen-II itself, via the C-propeptide domain, with Cys719 of another procollagen-II strand, or with a number of disulfide-forming proteins within the ER. Moreover, the Cys residue presents novel opportunities for redox-active components of the ER proteostasis network to recognize the presence of a misbehaving procollagen molecule.

Strikingly, despite Arg719Cys procollagen-II’s slow folding and induction of ER dilation, it is not detectably recognized as misbehaving by the ER proteostasis machinery. No component of the procollagen-II interactome engages Arg719Cys procollagen-II more than the wild-type protein (**Tables 1** and **S1**). Instead, the misbehaving procollagen appears to evade cellular quality control entirely. Consistent with this conclusion, Arg719Cys procollagen-II does not elicit any organized cellular stress response, such as the UPR (**Fig. 4**). In future work, it will be interesting to assess whether this behavior is recapitulated across other Arg➔Cys substitutions, notably those engendering much more severe pathologies.

While we did not observe any robust stress response in our transcriptional analysis, gene set enrichment analysis did reveal modest downregulation of translation-related gene sets (**Fig. S3**). Speculatively, this seemingly non-UPR-mediated transcriptional downregulation may reflect an adaptive response to procollagen-II accumulation or ER dilation. That being said, the mechanisms by which this accumulation is detected and transcriptional downregulation of some translation-associated genes is initiated are not clear, as they apparently do not originate with the UPR. Nevertheless, the most substantial signature in our transcriptional data was a delay in maturation and early hypertrophy at day 44 (**Fig. 4F**). This observation is consistent with the retardation of skeletal development noted in one of the transgenic mouse models of this disease,^21^ and may be associated with the mild spondyloepiphyseal dysplasia phenotype in patients that can be caused by slow skeletal development.^7^ Similarly, upregulation of *OGN* (**Fig. 1D**) could be a marker of the arthritic phenotype.^49, 50^

In fact, this experiment suggests that Arg719Cys is, from the perspective of the ER proteostasis network (at least to the sensitivity of our analysis), indistinguishable from wild-type procollagen-II. In patient tissue, Eyre *et al.* noted a reduced quantity of Arg719Cys procollagen-II in cartilage compared to the expected 50% in heterozygotes,^19^ which begs the question: if the ER proteostasis network is not recognizing Arg719Cys procollagen-II as problematic, what is? A key caveat of the quantitative interactome experiment is that, owing to the requirement for high-quality immunoprecipitation and a lack of suitable collagen-II antibodies, it was conducted by overexpressing epitope-tagged procollagen-II constructs in HT-1080 cells rather than in the more physiologically relevant chondronoids. Perhaps we were unable to detect some key ER proteostasis network factor. Other possible explanations for the reduced Arg719Cys procollagen-II content in patient tissue are post-ER quality control mechanisms recognizing the pathologic strand, and/or degradation mechanisms within the matrix selectively targeting the protein after secretion.^51^

Altogether, the novel human chondronoid model paired with an isogenic control reported and characterized here provides a robust and tractable platform for studying this disorder and a basis for modeling related Arg substitutions that drive other collagenopathies. Arg719Cys procollagen-II expression appears to cause an ER storage defect without inducing an associated ER stress response. Most significantly, it seems that the cell’s failure to effectively detect, much less resolve, the underlying protein folding challenge leads to the deposition of a defective collagen-II matrix. That defective matrix ultimately drives pathology. Thus, cellular strategies aiming to enhance detection and/or clearance of the problematic protein provide a novel therapeutic direction to explore.^3^ By virtue of its scalability, the chondronoid model reported here should prove valuable as a screening platform for interventions that correct or alleviate the disease phenotypes.

## METHODS

### General Reagents

All media and cell culture reagents were obtained from Corning/Cellgro, unless otherwise noted. Restriction enzymes, ligases, and polymerases were obtained from New England BioLabs. HA-Agarose beads (A2095) were obtained from Millipore. All cell lines used (iPSC lines and HT-1080) were regularly tested for mycoplasma and confirmed free of contamination.

Additionally, cells were continuously monitored for signs of bacterial contamination, doubling time, and general morphology.

### Plasmids

A plasmid encoding wild-type human *COL2A1* (variant IIB, consensus sequence) fused to GFP was obtained from Origene (#RG221644; NM_033150). The C-terminal GFP tag was removed and replaced by a stop codon. A pre-protrypsin signal sequence, HA epitope tag, and NotI site were inserted at the N-terminus of the N-propeptide using Gibson assembly. The Arg719Cys *COL2A1* plasmid was generated via site-directed mutagenesis of the wild-type plasmid. Both plasmids were sequence-confirmed via a full plasmid sequencing service offered by Plasmidosaurus.

### Antibodies

Antibodies were obtained from the following suppliers, and used at the specified dilutions: anti-Collagen-II (Millipore MAB8887; 1:150), anti-Calreticulin (Invitrogen AB_325990, PA3-900; 1:200), Goat Anti-Rabbit Alexa Fluor 488 (Abcam ab150077; 1:500), Goat Anti-Mouse Alexa Fluor 568 (Abcam ab175473; 1:500). The collagen-II antibody was validated on a histological section of a paraffin-embedded juvenile mouse knee (under 2 weeks of age; containing cartilage, ligament, and bone) to ensure proper selectivity for collagen-II. The calreticulin antibody has been widely used for immunohistochemistry, and was validated by the manufacturer on Western blot and on a section of rat brain cortex.

### Induced pluripotent stem cell lines growth and maintenance

Generating the gene-edited lines from ATCC fibroblast line 1502 was covered by the Human Research Ethics Committee of the Royal Children’s Hospital, Victoria, Australia (HREC #33118).^52^ The validation of iPSC lines MCRIi019-A (wild-type control) and MCRIi019-A-6 (*COL2A1* p.Arg719Cys heterozygote) was previously published.^25^ Briefly, it involved short tandem repeat profiling, karyotyping, and directed differentiation to all three germlines. iPSC lines were grown in a feeder-free manner on Matrigel (Corning)-coated 6-well plates in Essential 8 (E8) medium (ThermoFisher Scientific). Media was changed daily. Cells were routinely passaged (1:6 split) every 3–4 d and gently detached from plates using 0.5 mM EDTA in phosphate-buffered saline (PBS).

### Chondrocyte differentiation

Undifferentiated iPSCs at 70–90% confluency were dissociated using 0.5 mM EDTA in PBS and passaged into 6-well plates (∼2 × 10^5^ cells per well) pre-coated with Matrigel, and cultured for 48 h in E8 expansion medium. Differentiation to sclerotome was performed as described,^23, 53^ with APEL2 (StemCell Technologies) containing 5% Protein Free Hybridoma Medium (PFHM II; ThermoFisher Scientific). On day 0, medium was changed to anterior primitive streak-inducing medium containing 30 ng/ml Activin A (R&D Systems), 4 μM CHIR99021 (Tocris), 20 ng/mL FGF2 (PeproTech) and 100 nM PIK90 (Merck Millipore). After 24 h, this medium was replaced with medium containing 3 μM CHIR99021, 20 ng/mL FGF2, 1 μM A8301 (Tocris) and 0.25 μM LDN193189 (Cayman Chemical) to induce paraxial mesoderm for 24 h. Early somite development was then induced with medium containing 1 μM A8301 and 0.25 μM LDN193189, 3 μM C59 (Tocris) and 0.5 μM PD0325901 (Selleck Chemicals). After 24 h, sclerotome induction was initiated with 1 μM C59 and 2 μM purmorphamine (Sigma-Aldrich). After the first day of sclerotome induction (day 4), cells were dissociated from monolayer culture using Tryple Select (ThermoFisher Scientific), resuspended in sclerotome differentiation media, 300 μL aliquots containing 2 × 10^5^ cells were dispensed into 96-well low-attachment, round-bottom plates (Corning), and pelleted by centrifugation at 400 × *g* for 3 min using a swing-out rotor. Pellets were incubated in sclerotome differentiation media for a further 48 h to complete sclerotome differentiation in a 96-well static culture format. Differentiations were performed in biological triplicate, where individual replicates originated from cells of separate plates differentiated with individually prepared differentiation media from different media bottles.

### Chondrocyte maturation and extracellular matrix deposition

To promote chondrocyte differentiation, maturation, and ECM deposition, sclerotome cell pellets were cultured in 96-well round bottom plates in APEL2 medium supplemented with 5% PFHM II, 20 ng/mL FGF2 (PeproTech), 1× Penicillin-Streptomycin (ThermoFisher Scientific), and 200 μM freshly added sodium ascorbate (Amresco). On day 20, pellets were transferred to 6-cm non-adherent culture dishes (Greiner) with 15–20 pellets per dish in 5 mL of APEL2/PFHM II medium lacking FGF2 with orbital rotation at 60 rpm. Media was changed every 2–3 d for an additional 24 d of maturation. For harvesting, chondronoids were rinsed once with PBS and snap-frozen or processed for downstream assays. For each assay, chondronoids were harvested from all three biological replicates. Chondronoids originating from the same differentiation were considered technical replicates.

### Histology and Toluidine blue staining

Cultured chondronoids were fixed overnight at 4 °C in 10% neutral-buffered formalin (Millipore), washed in 70% ethanol, processed on a tissue processor (Sakura Tissue Tek VIP5) with a biopsy processing schedule as follows: 70% ethanol – 10 mins, 95% ethanol×2 – 10 mins each, 100% ethanol×3 – 10 mins each, xylene×2 – 10 mins each, Paraffin wax×4 – 10 mins each, and then embedded in “Paraplast X-tra” paraffin wax using a Leica embedding station. Serial 4 μm sections were cut from the chondronoids using a Leica rotary microtome. Sections were floated onto a 45 °C water bath and mounted on Leica “X-tra” adhesive slides (Leica Biosytems), drained, and dried at rt o/n. Before staining, sections were heated at 60 °C for 30 min. Sections were treated with xylene to remove the paraffin wax and then with an ethanol series comprising 100%, 90%, 70% ethanol, followed by water to rehydrate the sections.

Sections were stained with Toluidine blue solution (1% Toluidine blue, 0.2 M sodium phosphate at pH 9, filtered) for 10 min to detect a cartilage proteoglycan matrix, then rinsed in distilled water. Microscope slides were cover slipped using ProLong Gold Antifade Reagent (Invitrogen) and sealed with nail polish. Images were captured using a Leica DM 2000 LED microscope with Leica Application Suite (LAS) software version 4.9.0. Image processing was performed using ImageJ.

### Immunohistochemistry and confocal microscopy

Rehydrated histological sections underwent antigen retrieval by incubation with 1 mg/mL pepsin in 0.5 M acetic acid (Millipore) at 37 °C for 30 min. Samples were washed 3× with PBS, then incubated with 0.1% Triton X-100 for 30 min to permeabilize the cells. Samples were then incubated in a solution of 5% bovine serum albumin (BSA; Millipore) in PBS at rt for 1 h to block non-specific antibody binding. Samples were labeled with anti-Col-II antibody (Millipore MAB8887; 1:150) in 1% BSA for 1 h at rt. Samples were washed 3× with PBS. Alexa Fluor 568-conjugated anti-mouse secondary antibody was applied in 1% BSA solution for 30–60 min at rt. Samples were washed 3× with PBS. Cover slips were mounted on microscope slides using ProLong Gold Antifade Reagent with DAPI (Invitrogen) and sealed with nail polish. Images were acquired at the W. M. Keck Microscopy Facility at the Whitehead Institute on a Zeiss AxioVert200M microscope with a 10×, 63×, or 100× oil immersion objective with a Yokogawa CSU-22 spinning disk confocal head with a Borealis modification (Spectral Applied Research/Andor) and a Hamamatsu ORCA-ER CCD camera. The MetaMorph software package (Molecular Devices) was used for image acquisition. The excitation lasers used to capture the images were 405 nm, 488 nm, and 561 nm. Image processing was performed using ImageJ.

### TUNEL assay

Rehydrated histological sections were assayed for apoptosis using an *in situ* cell death detection kit (Millipore), according to the manufacturer’s instructions. At least four different chondronoids per genotype were used for analysis, with 2–3 sections of each. Briefly, following cell permeabilization, DNA breaks were labeled directly by the enzyme terminal deoxynucleotidyl transferase (TdT) with fluorescein labeled dUTPs. The positive control sample was incubated with DNAse for 15 min at 37 °C prior to TdT labeling. The negative control sample did not include the TdT enzyme. Samples were visualized on a fluorescence microscope (ECHO Revolve 4). Signal quantification was performed on the ECHO Pro App by analyzing the pixel intensity over the highlighted region of interest. Intensity of the multiple sections were averaged for each chondronoid, and significance was assessed using a homoscedastic student’s *t*-test.

### Transmission electron microscopy

Samples were washed with PBS and fixed with 2.5% glutaraldehyde, 2.0% *para*-formaldehyde in 100 mM cacodylate buffer pH 7.2 for 2 h at 4 °C, then post-fixed with 1% osmium tetroxide in 1.25% potassium ferrocyanide. They were then *en-bloc* stained with 2% uranyl acetate in 0.05 M maleate buffer pH 5.2 overnight at rt, followed by serial dehydrations with ethanol, then embedded in resin at 60 °C for 48 h. 60 nm sections were obtained using a diamond knife on a Leica UC67 Ultramicrotome and observed at 120 kV on a T12 Spirit Transmission Electron Microscope (ThermoFisher Instruments). Micrographs were captured using a digital camera from Advanced Microscopy Techniques (AMT).

### RNA-sequencing

For RNA-sequencing, individual chondronoids were snap-frozen in liquid nitrogen and pulverized using a liquid nitrogen-cooled tissue grinder. RNA was extracted with TRIzol (Invitrogen), followed by purification using Direct-zol RNA Microprep kit spin columns (Zymo Research), according to the manufacturer’s instructions. RNA samples were run on a fraction analyzer at MIT’s BioMicro Center for quality control. All RNA samples were processed at the same time using the same batch of reagents. RNA samples were quantified using an Advanced Analytical Fragment Analyzer. 10 ng of total RNA was used for library preparation on a Tecan ECO150. Custom primers were used to append a 6 bp barcode specific to each sample, as well as a unique molecular identifier (Integrated DNA technologies). Maxima H Minus Reverse Transcriptase (Thermo) was added per the manufacturer’s recommendations with a template-switching oligo, incubated at 42 °C for 90 min, and inactivated by incubation at 80 °C for 10 min. Following the template switching reaction, cDNA from 24 wells containing unique well identifiers were pooled together and cleaned using RNA Ampure beads at 1.0×. cDNA was eluted with 17 μL of water, digested with Exonuclease-I (New England Biolabs) at 37 °C for 30 min, and inactivated by incubation at 80 °C for 20 min. Second strand synthesis and PCR amplification was performed by adding Advantage 2 Polymerase Mix (Clontech) and the SINGV6 primer (Integrated DNA Technologies, 10 pmol, 5’-/5Biosg/ACACTCTTTCCCTACACGACGC-3’) directly to half of the exonuclease reaction volume. Eight cycles of PCR were performed, followed by clean-up using regular SPRI beads at 0.6× and elution with 20 μL of RSB. Successful amplification of cDNA was confirmed using the Fragment Analyzer. Illumina libraries were then produced using Nextera FLEX tagmentation substituting P5NEXTPT5-bmc primer (25 μM; Integrated DNA Technologies; 5’-AATGATACGGCGACCACCGAGATCTACACTCTTTCCCTACACGACGCTCTTCCG*A*T *C*T*-3’ where * = phosphorothioate bonds.) in place of the normal N500 primer. Final libraries were cleaned using SPRI beads at 0.7 × and quantified using the Fragment Analyzer and qPCR before being loaded for paired-end sequencing using an Illumina NextSeq500 in high-output paired-end mode (20/57 nt reads). Analyses were performed using previously described tools and methods. Reads were aligned against hg19 (February 2009) using BWA mem v.0.7.12-r1039 (RRID:SCR_010910) with flags –t 16 –f, and mapping rates, fraction of multiply-mapping reads, number of unique 20-mers at the 5’ end of the reads, insert size distributions and fraction of ribosomal RNAs were calculated using BEDTools v. 2.25.0 (RRID:SCR_006646).^54^ In addition, each resulting bam file was randomly down-sampled to a million reads, which were aligned against hg19, and read density across genomic features were estimated for RNA-Seq-specific quality control metrics. For mapping and quantitation, reads were scored against GRCh38/ENSEMBL 101 annotation using Salmon v.1.3 with flags quant -p 8 -l ISR – validateMappings.^55^ The resulting quant.sf files were imported into the R statistical environment using the tximport library (tximport function, option “salmon”), and gene-level counts and transcript per milllion (TPM) estimates were calculated for all genes.^56^ Samples were clustered based on log_2_-transformed TPMs of protein-coding genes and obvious outliers were excluded from downstream analyses. After filtering, each genotype group was made up of sequencing data from at least nine individual chondronoids.

RNA-Seq count data are not expected to follow a normal distribution, as their mean-variance relationship is not linear but instead follows a negative-binomial distribution. These data were modeled in the DESeq2 statistical framework, which entails a computation of the mean-variance relationship for each gene and its corollary dispersion. The underlying assumption of the algorithm is that the bulk of gene expression will remain unchanged among conditions. Using dispersion, statistically outlying genes were identified, and the significance of their putative differential expression was assessed between conditions.

Differential expression was also analyzed in the R statistical environment (R v.3.5.1) using Bioconductor’s DESeq2 package (RRID:SCR_000154).^57^ Dataset parameters were estimated using the estimateSizeFactors(), and estimateDispersions() functions; read counts across conditions were modeled based on a negative binomial distribution, and a Wald test was used to test for differential expression (nbinomWaldtest(), all packaged into the DESeq() function), using the treatment type as a contrast. Shrunken log_2_ fold-changes were calculated using the lfcShrink function, based on a normal shrinkage estimator.^57^ Fold-changes, *p*-values and false-discovery rates were reported for each gene. All the gene sets considered were either previously published (UPR,^58^ non-canonical stress,^43, 59^ ISR,^44^ early hypertrophy^23^), or curated from previously described gene ontologies.

### Gene Set Enrichment Analysis (GSEA)

Differential expression results from DESeq2 were retrieved, and the “stat” column was used to pre-rank genes for GSEA analysis. These “stat” values reflect the Wald’s test performed on read counts as modeled by DESeq2 using the negative binomial distribution. Genes that were not expressed were excluded from the analysis. GSEA (linux desktop version, v4.1)^60^ was run in the pre-ranked mode against MSigDB 7.0 C5 (Gene Ontology) set, and ENSEMBL IDs were collapsed to gene symbols using the Human_ENSEMBL_Gene_ID_MSigDB.v7.4.chip, resulting in 25162 and 26199 genes for wild-type versus mutant sample in early and late time-point comparisons, respectively. In addition, a weighted scoring scheme, meandiv normalization, cutoffs on MSigDB signatures sizes (between 5 and 2000 genes, resulting in 9,476 and 9,588 gene sets retained for early and late time-point wild-type vs. mutant comparisons, respectively) were applied and 5,000 permutations were run for *p*-value estimation.

### Collagen extraction and SDS-PAGE analysis

Chondronoids of the same genotype were rinsed with PBS, blotted dry with a Kimwipe, pooled, snap frozen in liquid nitrogen, and pulverized using a liquid nitrogen-cooled tissue grinder. The powder obtained was added to 100 μL chondroitinase ABC solution (50 mM Tris HCl pH 8, 50 mM sodium acetate, 0.1 mU/μL chondroitinase ABC (Millipore)) and incubated in a thermomixer set to 37 °C and 1,500 rpm for 6 h. 900 μL of denaturing solution (50 mM sodium acetate, 10 mM EDTA, 4 M guanidine hydrochloride, 65 mM dithiothreitol) was added, and samples were rotated end-over-end at 4 °C overnight. The next day, samples were centrifuged at 13,000 rpm at 4 °C for 10 min. The supernatant was removed and the pellet resuspended in 200 μL of cold pepsin solution [0.1 mg/mL pepsin (Millipore) in 0.5 M acetic acid] and rotated end-over-end at 4 °C o/n. The next day, samples were centrifuged at 13,000 rpm at 4 °C for 10 min. The supernatant was saved, neutralized with 10 M NaOH, and lyophilized. Lyophilized samples were reconstituted in loading buffer for SDS-PAGE [50 mM Tris HCl pH 6.8, 2 M urea, 0.1% SDS, 0.1% sucrose, 0.01% bromophenol blue (Millipore)] and boiled for 15 min. Samples were separated on homemade 4/8% tris SDS-PAGE gels. Gels were stained in Coomassie Blue solution [0.1% Coomassie Blue (Millipore), 10% acetic acid, 50% methanol, 40% water] and destained in the same solution without Coomassie Blue. Gels were imaged by placing them on a lightbox and photographing them with a smartphone. A raw image of the full gel is provided in the SI.

### Intracellular collagen-II quantification

A survey of confocal images of immunohistochemistry samples stained for collagen-II was used to quantify the percent of cells with intracellularly retained collagen-II. 15–20 images of the different genotypes at different magnifications (40–100×) were arranged in a random order. Each genotype set consisted of at least three individual chondronoids. Three independent and blinded participants were asked to count the total number of cells and the number of cells with intracellular staining in each image. The proportion of positive cells was averaged between all participants for each image. A total of >120 cells were analyzed per genotype. Statistical significance between genotypes was calculated using an unpaired homoscedastic student’s *t*-test, where each image represents a replicate.

## Proteomics

### Cell Culture, transfections, and immunoprecipitation

HT-1080 cells (ATCC) were cultured in complete DMEM supplemented with 10% FBS. Cells from ATCC were used without further authentication. Transfections of wild-type and Arg719Cys COL2A1-encoding plasmids were performed using Transit-2020 transfection reagent (Mirus), according to the manufacturer’s instructions. The next day, media was replaced with fresh complete DMEM supplemented with 50 μM sodium ascorbate (Amresco). Two days post-transfection, cells were harvested, washed with 1× PBS, and crosslinked in the plate using 0.2 mM dithiobis(succinimidyl propionate) (DSP; Lomant’s Reagent, Thermo) in PBS at rt for 30 min. Crosslinking was quenched by addition of 0.1 M Tris at pH 8.0 (final concentration) for 10 min at rt. Cells were scraped from plates in PBS, then pelleted and resuspended in RIPA [150 mM sodium chloride, 50 mM Tris-HCl, pH 7.5, 1% Triton X-100, 0.5% sodium deoxycholate, 0.1% sodium dodecyl sulfate (SDS), protease inhibitor tablets and 1.5 mM phenylmethylsulfonyl fluoride (PMSF; Amresco)]. Cells were lysed for 30 min on ice, then centrifuged at 21,100 × g for 15 min at 4 °C. Supernatants were collected, quantified for protein content (BCA assay; Pierce), normalized, and incubated for 16 h with HA antibody-conjugated beads, rotating end-over-end at 4 °C. Samples were washed with RIPA 3× and then eluted by boiling for 10 min in 300 mM Tris at pH 6.8 with 6% SDS and 600 mM dithiothreitol (Millipore) to release crosslinks. Eluates were prepared for mass spectrometry analysis.

### Mass spectrometry and interactome characterization

Samples were reduced, alkylated, and digested with sequencing-grade trypsin (Promega) using an S-Trap Micro (Protifi) according to manufacturer’s instructions. After digestion, the samples were labeled with TMT 10plex™ reagents (Thermo), according to the manufacturer’s instructions. Briefly, the dried tryptic peptides were resuspended in 100 mM triethylammmonium bicarbonate (TEAB) and then vortexed. Acetonitrile was added to each of the 10-plex TMT label reagents, and then samples were vortexed and briefly centrifuged. Peptides and labeling reagents were then combined in a 1:2 ratio and incubated for 1 h at rt, after which the reaction was quenched with 5% hydroxylamine for 15 min. Equal amounts of each channel were combined, dried by speed-vacuum, and resuspended in TEAB. Samples were then fractionated with Pierce High pH Reversed-Phase peptide Fractionation Kits (catalog #84868) using standard protocols, then dried and resuspended in 0.2% formic acid for injection on LC-MS/MS. Labeled tryptic peptides were separated by reverse phase HPLC (Thermo Ultimate 3000) using a Thermo PepMap RSLC C_18_ column (2 μm tip, 75 μm × 50 cm, PN# ES903) over a gradient before nano-electrospray injection onto an Orbitrap Exploris 480 mass spectrometer (Thermo).

The mass spectrometer was operated in a data-dependent mode. The parameters for the full scan MS were: resolution of 120,000 across 375–1600 m/z and maximum IT of 25 ms. The full MS scan was followed by MS/MS for as many precursor ions in a three second cycle with a NCE of 32, dynamic exclusion of 30 s and resolution of 45,000. Raw mass spectral data files (.raw) were searched using Sequest HT in Proteome Discoverer (Thermo). Sequest search parameters were: 10 ppm mass tolerance for precursor ions; 0.05 Da for fragment ion mass tolerance; 2 missed cleavages of trypsin; fixed modification were carbamidomethylation of cysteine and TMT modification on the lysines and peptide N-termini; variable modifications were methionine oxidation, methionine loss at the N-terminus of the protein, acetylation of the N-terminus of the protein, methylation of histidine, arginine, and glutamine, hydroxylation of proline, and methionine-loss plus acetylation of the protein N-terminus; total PTM per peptide was set to 8. The false discovery rate was set at 0.01. Data were searched against a human database and a common MS contaminant database made in-house. Human Database: www.uniprot.org/proteomes/UP000005640.

Raw abundance values were used for quantification. First, abundance of each protein in the negative control was determined as the average abundance of the four no-transfection control conditions. High confidence interactors were identified as any protein whose TMT intensities were highly correlated with those of the bait in wild-type samples,^61^ as assessed by a Pearson’s correlation q-value < 0.02. For each experimental sample, the abundance of each protein was normalized to the abundance of bait (procollagen-II) to obtain a normalized abundance value.

The fold-enrichment was calculated as the ratio of the average of normalized abundance values in the Arg719Cys procollagen-II IP to the average of normalized abundance values in the wild-type procollagen-II IP.^61^ Significance was evaluated using an unpaired homoscedastic student’s t-test across the four biological replicates, and corrected for multiple hypothesis testing using a two-stage step-up method of Benjamini, Krieger, and Yekutieli with a desired FDR set to 1%.

## ACKNOWLEDGEMENTS

The authors would like to acknowledge Dr. Louise Kung, Lisa Sampurno, and Lynn Rowley, and Jinia Lilianty at Murdoch Children’s Research Institute for generously sharing their expertise and technical knowledge in iPSC culture, differentiation, and analysis. We thank Margaret Bisher, David Mankus, and Dr. Abigail Lytton-Jean at the Koch Institute for Integrative Cancer Research’s Robert A. Swanson Biotechnology Center Nanotechnology Materials Facility for their expertise and technical assistance with electron microscopy, as well as Caroline Atkinson at the MIT Division of Comparative Medicine for her valuable technical support and advice. We extend our gratitude to Brandyn Braswell and others at the W.M. Keck Microscopy Facility at the Whitehead Institute for technical support with confocal microscopy, and to Dr. Stuart Levine at the MIT BioMicro Center for assistance with RNA-sequencing. We thank Dr. Jimin Yoon, Jessica Patrick, and Sorin Srinivasa for generously offering their time to quantify intracellular collagen. This work was supported by a Research Grant from the G. Harold and Leila Y. Mathers Foundation (to M.D.S., S.R.L., and J.F.B.), the National Institutes of Health (NIH; AR082995 to M.D.S.), the Australia National Health & Medical Research Council (GNT2003393 and GNT1146952 to both S.R.L. and J.F.B., and GNT1146902 to J.F.B.), and the Victorian Government’s Operational Infrastructure Support Program. K.M.Y. was supported by an NIH Ruth L. Kirschstein Predoctoral Fellowship (F31AR079263). S.K. was supported by a Kwanjeong Fellowship. This work was also supported in part by the National Cancer Institute (Koch Institute core) Grant P30-CA14051 and the National Institute of Environmental Health Sciences (core) Grant P30-ES002109. Work in the Novo Nordisk Foundation Center for Stem Cell Medicine is supported by a Novo Nordisk Foundation grant (NNF21CC0073729).

