## Supplemental Information for "Failed Cellular Surveillance Enables Pathogenic Matrix Deposition in a *COL2A1*-Related Osteoarthritis"

| **Page** | **Content** |
| --- | --- |
| S1 | Table of Contents |
| S2–3 | Supporting Table |
| S4 | Supporting Figure Legends |
| S5–8 | Supporting Figures |

**SUPPORTING TABLE**

| **Table S1.** High-confidence interactors*^a^* engage similarly with wild-type and Arg719Cys procollagen-II across four biological replicates. | | | |
| --- | --- | --- | --- |
| **Protein (*Gene*)** | **Fold enrichment*^b^*** | ***p*-value*^c^*** | ***q*-value*^d^*** |
| Galectin-3-binding protein (*LGALS3BP*) | 0.71 | 0.06 | 0.33 |
| DnaJ homolog subfamily B member 11 (*DNAJB11*) | 0.81 | 0.01 | 0.23 |
| ADP/ATP translocase 2 (*SLC25A5*) | 0.82 | 0.28 | 0.50 |
| Fibronectin (*FN1*) | 0.83 | 0.21 | 0.47 |
| Protein-glutamine gamma-glutamyltransferase 2 (*TGM2*) | 0.84 | 0.36 | 0.53 |
| Endoplasmic reticulum chaperone BiP (*HSPA5*) | 0.86 | 0.07 | 0.33 |
| Prohibitin (*PHB2*) | 0.90 | 0.67 | 0.72 |
| Protein disulfide-isomerase A6 (*PDIA6*) | 0.94 | 0.47 | 0.61 |
| Heat shock protein 90 beta family member 1 (*HSP90B1*) | 0.94 | 0.25 | 0.50 |
| Calumenin (*CALU*) | 0.96 | 0.64 | 0.74 |
| Calreticulin (*CALR*) | 0.98 | 0.86 | 0.89 |
| RAB1B, member RAS oncogene family (*RAB1B*) | 0.99 | 0.97 | 0.98 |
| Inhibitor of nuclear factor kappa-B kinase-interacting protein (*IKBIP*) | 0.99 | 0.91 | 0.89 |
| Prolyl 4-hydroxylase subunit alpha-2 (*P4HA2*) | 1.03 | 0.57 | 0.70 |
| Neutral alpha-glucosidase AB (*GANAB*) | 1.04 | 0.62 | 0.72 |
| Serpin H1 [Hsp47] (*SERPINH1*) | 1.06 | 0.07 | 0.33 |
| Prolyl 4-hydroxylase subunit alpha-1 (*P4HA1*) | 1.06 | 0.27 | 0.50 |
| Multifunctional procollagen lysine hydroxylase and glycosyltransferase LH3 (*PLOD3*) | 1.07 | 0.21 | 0.47 |
| Reticulocalbin-1 (*RCN1*) | 1.07 | 0.27 | 0.50 |
| Glucosidase 2 subunit beta (*PRKCSH*) | 1.08 | 0.46 | 0.61 |
| Calnexin (*CANX*) | 1.09 | 0.76 | 0.83 |
| Prolyl 3-hydroxylase 1 (*P3H1*) | 1.10 | 0.16 | 0.47 |
| Protein disulfide-isomerase A3 (*PDIA3*) | 1.11 | 0.16 | 0.47 |
| Procollagen galactosyltransferase 1 (*COLGALT1*) | 1.11 | 0.35 | 0.53 |
| Dolichyl-diphosphooligosaccharide-protein glycosyltransferase subunit 1 (*RPN1*) | 1.11 | 0.63 | 0.73 |
| DnaJ homolog subfamily C member 3 (*DNAJC3*) | 1.12 | 0.50 | 0.53 |
| Peptidyl-prolyl cis-trans isomerase FKBP10 (*FKBP10*) | 1.12 | 0.11 | 0.43 |
| Protein disulfide-isomerase (*P4HB*) | 1.13 | 0.15 | 0.47 |
| Protein disulfide-isomerase A4 (*PDIA4*) | 1.15 | 0.23 | 0.50 |
| Cytoskeleton-associated protein 4 (*CKAP4*) | 1.16 | 0.08 | 0.35 |
| Dolichyl-diphosphooligosaccharide-protein glycosyltransferase subunit 2 (*RPN2*) | 1.17 | 0.50 | 0.61 |
| Erlin-2 (*ERLIN2*) | 1.17 | 0.21 | 0.47 |
| Procollagen-lysine,2-oxoglutarate 5-dioxygenase 1 (*PLOD1*) | 1.17 | 0.30 | 0.53 |
| Hypoxia upregulated 1 (*HYOU1*) | 1.20 | 0.19 | 0.47 |
| Cartilage-associated protein (*CRTAP*) | 1.24 | 0.02 | 0.22 |
| Peptidyl-prolyl cis-trans isomerase FKBP9 (*FKBP9*) | 1.28 | 0.04 | 0.33 |
| Peptidyl-prolyl cis-trans isomerase (*PPIB*) | 1.38 | 0.03 | 0.33 |
| Procollagen-lysine 5-dioxygenase (*PLOD2*) | 1.39 | 0.01 | 0.22 |
| Thioredoxin domain-containing protein 5 (*TXNDC5*) | 1.54 | 0.06 | 0.33 |
| Thioredoxin (*TXN*) | 1.86 | 0.33 | 0.53 |
| Triosephosphate isomerase (*TPI1*) | 3.43 | 0.38 | 0.53 |
| Protein S100-A11 (*S100A11*) | 6.21 | 0.37 | 0.53 |
| *^a^*High confidence interactors were identified as any protein whose TMT intensities were highly correlated with those of the bait in wild-type samples, as assessed by a Pearson’s correlation *q* value < 0.02.  *^b^*Fold-enrichment was calculated as the ratio of bait-normalized abundance of peptides detected in Arg719Cys to the bait-normalized abundance of peptides detected in wild-type.  *^c^p*-value was calculated using a homoscedastic student’s *t*-test across the four biological replicates.  *^d^q*-value was calculated using a two-stage step-up method of Benjamini, Krieger, and Yekutieli with a desired false discovery rate set to 1%. | | | |

**SUPPORTING FIGURE LEGENDS**

**Figure S1** Full SDS-PAGE image. Chondronoid pepsin-extracted collagen was run on a homemade 4/8% SDS-PAGE gel, and then stained with Coomassie blue dye.

**Figure S2** Interactomic workflow to identify differential interactors. Cells were transfected with a plasmid expressing either wild-type or Arg719Cys procollagen-II fused to an N-terminal HA epitope tag, in biological quadruplicate. The negative control consisted of non-transfected cells (no procollagen-II expressed). Two days post-transfection, proteins were crosslinked using Lomant’s cell permeable crosslinking reagent, then lysed. Procollagen-II and its interactors were immunoprecipitated with anti-HA agarose beads, reduced to release the crosslinks, and processed for MS. TMTs were used to label peptides from each sample. All tagged peptides from a given replicate were then pooled and injected together for quantitative MS.

**Figure S3** GO pathways differentially expressed in Arg719Cys chondronoids compared to wild-type chondronoids at day 34. Pathways shown have a false discovery rate (FDR) < 1%. No pathways were significantly differentially expressed (FDR < 10%) at day 44. Differential expression was established using a preranked Wald’s test results from differential expression testing in DESeq2. Empirical significance of differential pathway representation was estimated using 5000 gene set permutations and FDRs were calculated using the Benjamini-Hochberg procedure. Genes that were not expressed were excluded from the analysis.

**Figure S4** TUNEL assay and controls. **(A)** Representative images of wild-type and Arg719Cys chondronoids stained with TUNEL. **(B)** Quantification of the signal across at least 4 individual chondronoids revealed no statistically significant difference in the extent of apoptosis between genotypes. Significance was evaluated using an unpaired, two-tailed *t* test. **(C)** The positive control sample was incubated with DNAse for 15 min at 37 °C to induce DNA strand breaks. The negative control sample was incubated with labeled nucleotides in the absence of the TdT enzyme. The high exposure image is provided to show the sample.

**SUPPORTING FIGURES**

**FIGURE S1**

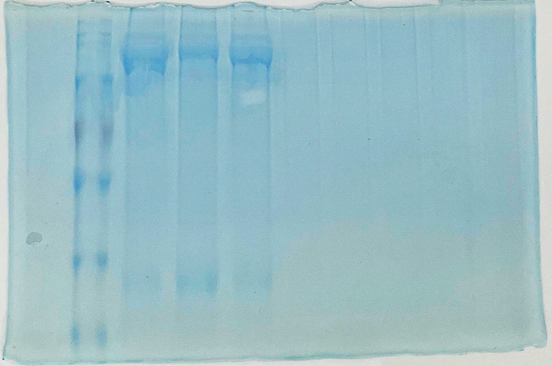

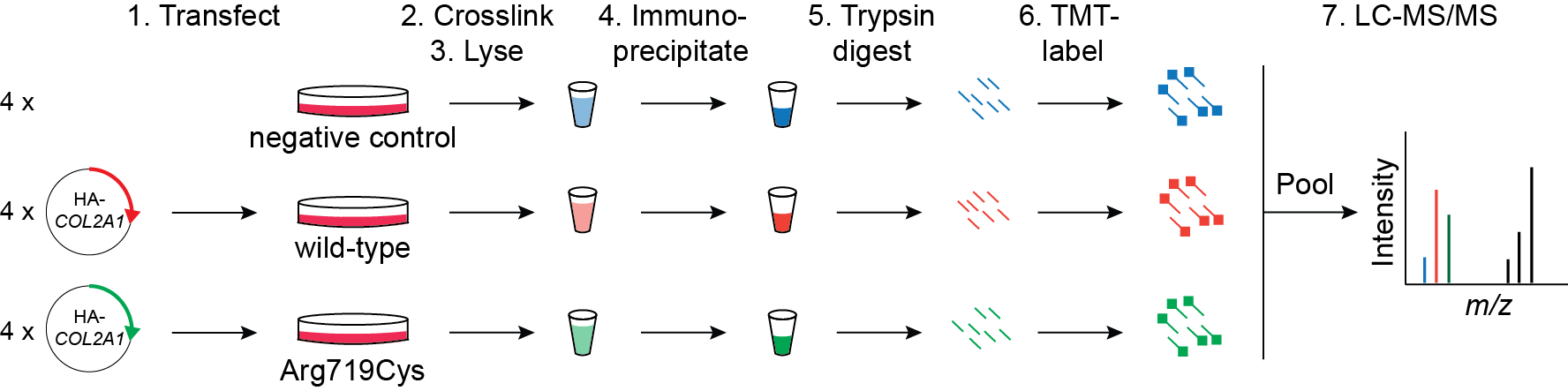
**FIGURE S2**

**FIGURE S3**

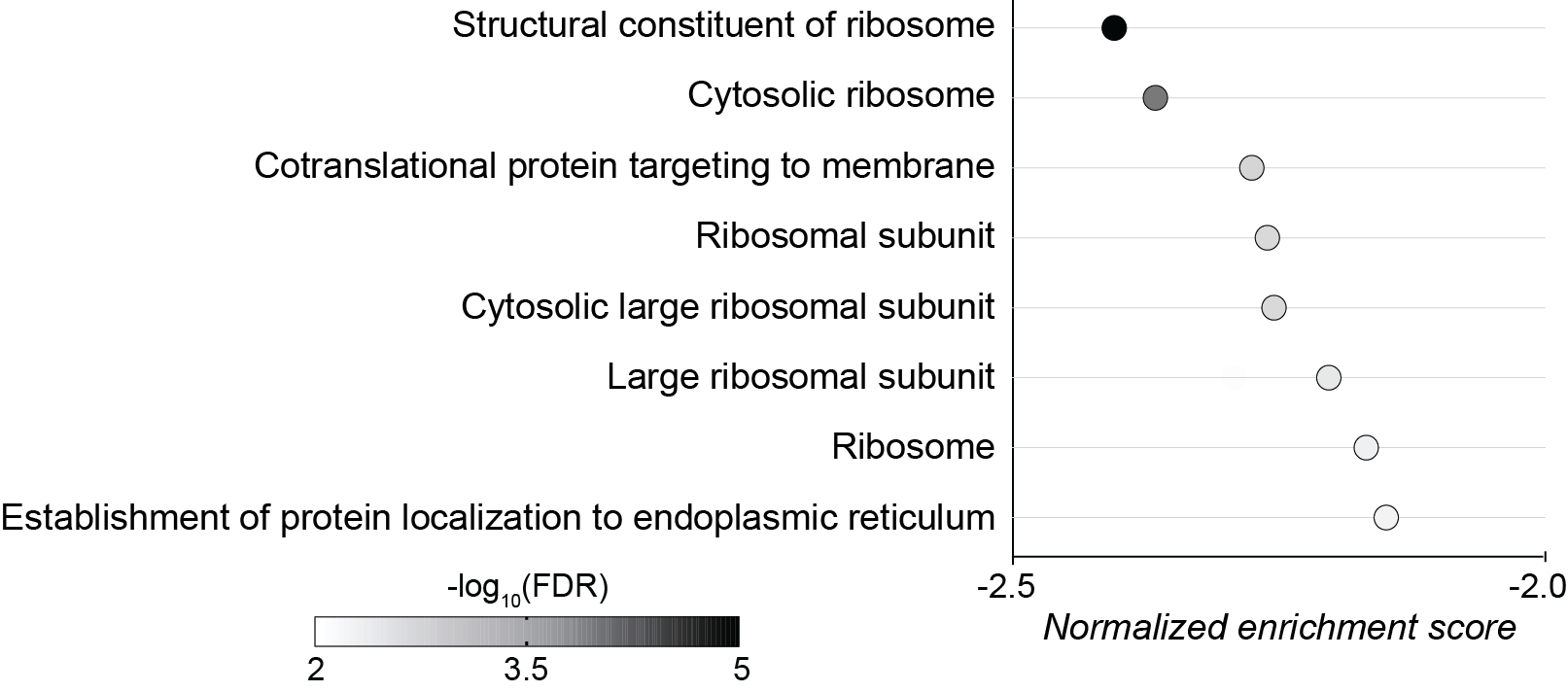

**FIGURE S4**

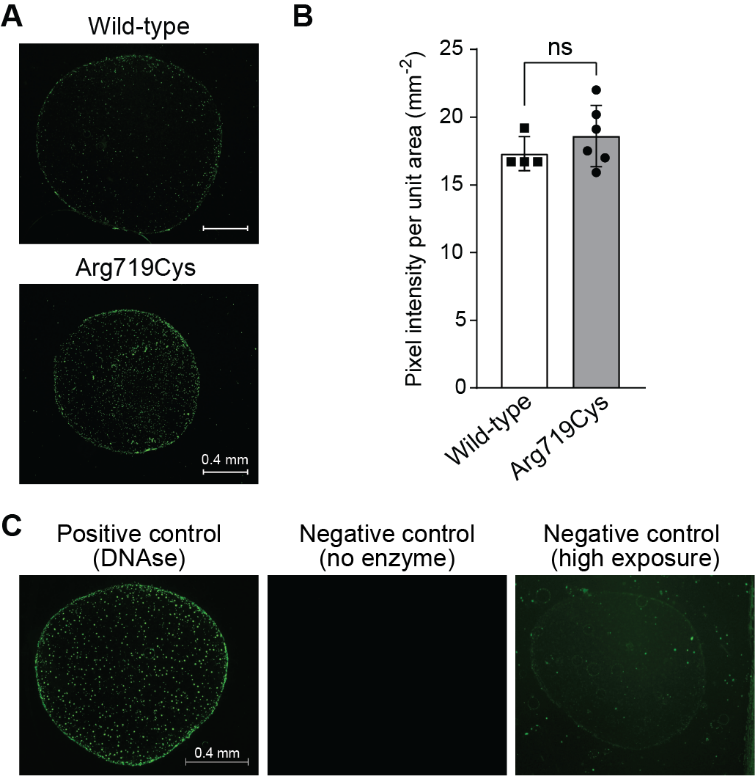
